# Cell-Free DNA Sequencing Uncovers the Longitudinal Consequences of Temozolomide Treatment and Host Co-Culture in Glioblastoma

**DOI:** 10.1101/2025.07.12.662613

**Authors:** Sharvari Mankame, Hersh Nanda, Maria Kyriakidou, Mimi Mbegbu, Nanyun Tang, Michael Berens, Floris Barthel

## Abstract

Glioblastoma (GBM) is a highly aggressive brain tumor with limited options for longitudinal monitoring. We evaluated the potential of cell-free DNA (cfDNA) as a real-time biomarker of tumor burden under tightly controlled conditions. We measured the cfDNA release dynamics, fragmentation sizes, and variant allele frequencies (VAF) in patient-derived GBM cell cultures. Time series of cfDNA were collected and analyzed in treatment-naïve monocultures, during temozolomide (TMZ) treatment, and in co-culture with normal human astrocyte (NHA) cells. Longitudinal collection of media from individual cultures demonstrated that cfDNA yield increased, indicating the ability to use cfDNA to track tumor burden over time. Using a co-culture system, we deconvoluted cfDNA admixtures by analyzing yield, fragment size patterns, and cell line–specific variants. The exclusive detection of NHA- and GBM-specific mutations confirmed the distinct contributions of each cell type. Finally, TMZ treatment of GBM cells prompted an increase in cfDNA yield and VAF, suggesting that the effects of therapy could be measured using cfDNA. These findings support cfDNA as a non-invasive biomarker for real-time monitoring of GBM progression and treatment response, with clinical potential as a liquid biopsy tool in glioblastoma management.

**KEY POINTS:**

1. cfDNA yield and variant allele frequencies (VAFs) correlate with GBM tumor burden.
2. cfDNA properties can be used to differentiate between cell types in a mixed population.
3. TMZ treatment increases cfDNA yield and VAFs, reflecting treatment response.

**IMPORTANCE OF THE STUDY:** This study highlights the potential of cell-free DNA (cfDNA) as a non-invasive biomarker for monitoring glioblastoma (GBM) progression and treatment response. By utilizing in-vitro models, we demonstrate that cfDNA yield, fragmentation patterns, and variant allele frequencies (VAFs) can effectively identify treatment-induced changes, such as those induced by temozolomide (TMZ) treatment, providing insights into therapeutic response. These results have significant translational implications, as cfDNA could serve as a real-time liquid biopsy tool for monitoring GBM progression, assessing treatment efficacy, and identifying early signs of treatment resistance in clinical settings. The ability to track tumor dynamics non-invasively holds great promise for improving GBM patient management and guiding personalized therapeutic approaches.

## INTRODUCTION

Glioblastoma multiforme (GBM), one of the deadliest forms of primary brain cancer, is characterized by rapid growth, diffuse infiltration, and aggressive vascular proliferation; all contributing to poor prognosis in patients. The current standard of care includes maximal safe surgical resection followed by radiotherapy and temozolomide (TMZ) chemotherapy. However, GBM’s ability to develop resistance to radiation and alkylating agents leads to an abbreviated survival benefit. Despite extensive research aimed at identifying molecular targets to slow glioma progression, the biological processes driving treatment resistance remain poorly understood.

In IDH wild-type GBM, methylation of the O-6-Methylguanine-DNA Methyltransferase (MGMT) promoter silences the gene, reducing the expression of the DNA repair protein that would otherwise counteract TMZ-induced damage^15^. As such, MGMT promoter methylation status has become the most common biomarker for predicting TMZ responsiveness. However, determining methylation status requires invasive tissue biopsies and, even when obtained, offers no real-time insight into treatment effectiveness. This underscores the urgent need for non-invasive methods to monitor therapeutic response as it unfolds.

Cell-free DNA (cfDNA) has emerged as a promising, non-invasive tool for molecular profiling and disease monitoring. Derived from both normal and tumor cells in cancer patients, cfDNA reflects the genetic landscape of malignancies and has been successfully used to track tumor burden and treatment response in breast, lung, and ovarian cancers^4,5^. However, its potential in GBM for capturing clonal evolution and resistance mechanisms remains underexplored.

Understanding the dynamics of TMZ resistance could pave the way for more personalized treatment strategies. In this study, we explore cfDNA as a liquid biopsy tool to monitor tumor burden in real-time, identify tumor-specific mutational signatures in various in-vitro models, and track the emergence of TMZ resistance in GBM by analyzing cfDNA yield, fragmentation patterns, and variant allele frequencies (VAFs) over time. This approach not only provides insights into glioma biology and the evolution of drug-resistant clones but also highlights cfDNA’s utility in various experimental paradigms.

By leveraging cfDNA’s non-invasive nature and temporal resolution, our work aims to enhance the management of GBM and ultimately improve patient outcomes.

## METHODS

### Cell Culture for GBM43

Patient-derived xenograft (PDX) GBM43 was derived from a male patient with a classical IDH-wildtype glioblastoma tumor. This cell line was maintained according to American Type Culture Collection recommendations at 37°C, 95% O_2_ and 5% CO_2_, in 1:1 parts of Dulbecco’s Modified Eagle Medium (DMEM) and F12 media supplemented with 5X B-27, 10X N-2, epidermal growth factor (EGF) and fibroblast growth factor (FGF). Cultures of this PDX exhibited adherent properties in standard adherent flasks. Prior to experimentation, the cells were maintained for several weeks with frequent passages when 80% confluency was reached. Once consistent viability (∼90%) was reached for consecutive passages, the cells were counted with the Countess Automated Cell Counter and 0.5E6 cells each were transferred into ten individual T75 flasks. The seeding density for the experiment was optimized based on the rate of confluency. Out of the ten flasks, five were labeled for the TMZ-treatment and five were labeled for vehicle (untreated) treatment. Following cell attachment after seeding (∼24 h), vehicle or treatment was applied. The TMZ dose was determined based on the IC50 value for PDC. For each treated flask receiving TMZ treatment, an equivalent volume of vehicle (media) was added to the corresponding untreated flask. Cell counts and supernatants were collected every 24 hours from each paired treated or untreated flask. All experiments were repeated in triplicate.

Upon detaching the cells from the flask, cells were spun down into a pellet and the supernatant was transferred into a different tube on ice. The supernatant underwent three more spins at 600rcf, 1500rcf and 6000rcf in a temperature-controlled centrifuge at 4C for 10 minutes each to minimize gDNA contamination. The cell pellet left behind from every spin step was combined with the original stock of cells from that time point for counting. In adherent cell cultures, dead cells are known to naturally detach and float in the supernatant and therefore the cells collected from each spin step consisted primarily of dead cells and were used to calculate the total dead cell count at each timepoint.

### Cell Culture for GBM12

PDX cell line GBM12 was collected from a male patient with an IDH-wildtype glioblastoma. This cell line was maintained according to American Type Culture Collection recommendations at 37°C, 95% O_2_ and 5% CO_2_, in 1:1 parts of Dulbecco’s Modified Eagle Medium (DMEM) and F12 media supplemented with 5X B-27, 10X N-2, epidermal growth factor (EGF) and fibroblast growth factor (FGF). Compared to GBM43, this cell line was grown as neurospheres and maintained at a similar timeline of passaging. The time course experiment was set up in the same way as GBM43 where cells were grown out and then divided amongst five flasks in equal 3E6 cell seeding densities. At each collection, the supernatant containing both the neurospheres and the cfDNA was collected and spun down at 1300rpm for 5 minutes. The supernatant was transferred into a new tube on ice and the remaining cell pellet was used to quantify the ratio of live and dead cells at each time point. The supernatant underwent the same spin steps as the GBM43 cultures in a 4C centrifuge for 10 minutes each to decrease gDNA carryover. This time course experiment was repeated in several patient-derived xenograft neurosphere cell lines (**Supplemental Figure 3**) and GBM12 was chosen for sequencing based on successful library preparation.

### Co-Culture with NHA and GBM43 cells

GBM43, an adherent cell line, was co-cultured with normal human astrocytes (NHA), also an adherent cell line at a 1:1 ratio. NHA cells were also individually grown out for the same five timepoints to assess growth rates and viability over time. GBM43 cells were tagged with RFP and NHA cells were dyed with GFP using CellTrace SYBR dye. In addition to cell counts using the automated Countess, the ratio of cell types was validated using fluorescent intensity.

### cfDNA extraction

The cfDNA in the purified supernatant was extracted using the MagMax cfDNA Isolation Kit according to the manufacturer’s instructions. The eluted cfDNA was quantified using Qubit dsDNA High Sensitivity (ThermoFisher, Waltham, MA, USA) and Agilent Tapestation High Sensitivity D1000 (Agilent, Santa Clara, CA, USA).

### cfDNA library preparation

Due to the low concentration of cfDNA, particularly in the untreated samples, we used 1 ng of cfDNA input to generate libraries using the Watchmaker DNA Library Preparation kit v2 with fragmentation, optimized for low input. 250bp fragments were generated following manufacturer’s protocol. Unique molecular indices (UMIs) were ligated to each sample at 20°C for 15 min. A 0.8X bead clean-up step using SPRI beads was performed prior to library amplification to purify each library. PCR was conducted using the following parameters: initial denaturation at 98°C for 45 sec, 11 cycles of: denaturation at 98°C for 15 sec, annealing at 60°C for 30 sec, extension at 72°C for 30 sec, and a final elongation at 72°C for 1 min. The PCR-amplified cfDNA libraries were purified again and then cleaned up using a 1X SPRI beads-to-sample ratio and eluted in 22ul of elution buffer. Library concentration and fragment size distribution were quantified with Qubit (ThermoFisher, Waltham, MA, USA) and Agilent Tapestation HS D1000 (Agilent, Santa Clara, CA, USA).

### gDNA extraction and library preparation

A cell pellet of 1E6 cells was taken for gDNA extraction from the same passage that was used to initiate the experiment for each of the three cell cultures: GBM43, GBM12 and NHA. The gDNA was extracted with QiAMP DNA Micro Kit using manufacturer recommended protocols. The gDNA was quantified using Qubit 1x dsDNA Broad Range (ThermoFisher, Waltham, MA, USA) and Agilent Tapestation gDNA (Agilent, Santa Clara, CA, USA). An indexed library for Illumina sequencing was prepared using the Twist EF 2.0 Kit (Twist Biosciences), which employed enzymatic fragmentation followed by end-repair, A-tailing, and PCR amplification. Starting with 500 ng of input DNA, samples were enzymatically sheared to an average fragment size of 250 bp and subjected to four cycles of PCR amplification.

### Target Enrichment

Target enrichment was performed using Twist Target Enrichment Fast Hybridization Protocol (Twist Biosciences). Each pool containing up to 8 libraries underwent target enrichment using a mitochondrial panel and a custom-made glioma panel consisting of highly mutated genes in glioblastoma (Twist Biosciences). Briefly, the libraries were pooled at equal concentrations to generate a 1.5ug pool and dried down using a vacuum centrifuge. Next, the pools are hybridized with the panels and incubated at 60°C for 4 hours. Next, streptavidin binding beads were used to capture the enriched DNA libraries. The captured libraries were washed with wash buffers and eluted in 45ul of molecular grade water. Each pool was then PCR amplified using the following parameters: initial denaturation at 98°C for 45 sec, 11 cycles of: denaturation at 98°C for 15 sec, annealing at 60°C for 30 sec, elongation at 72°C for 30 sec, and a final elongation at 72°C for 1 min. PCR products were purified using 1.8X DNA Purification beads (Twist Biosciences) and eluted in a final volume of 30ul. All pooled reactions were quantified using Qubit and Tapestation.

### Sequencing

All cfDNA and gDNA target enriched pools were run on the Illumina Novaseq X platform using 2×150 paired end sequencing. Due to size disparities amongst libraries, all gDNA pools were run separately from cfDNA pools.

### Computational Analysis

Due to the specialized UMI-UDI library preparation for all cfDNA and gDNA libraries, all reads utilized the fgbio pipeline optimized for UMI containing reads. First, empty reads were removed from the raw sequenced reads using cutadapt (v.4.8). All reads were mapped to the hg38 genome and PCR duplicates were identified and condensed into consensus sequences using BWA (v0.7.18) and fgbio (v 2.2.1). Somatic variants were called on the subsequent base recalibrated bam files using GATK’s mutect2. Low quality and germline variants were filtered out and the remaining variants were annotated using GATK Funcotator. The average coverage for each library was around 600x.

### Statistical Analysis

To evaluate changes in variant allele frequency (VAF) over time, we employed a linear mixed-effects model using the lme4 package in R (v4.0.2). The model was specified as

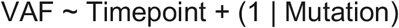

where Timepoint was treated as a fixed effect and Mutation (representing individual mutations) was modeled as a random intercept to account for repeated measures across mutations. To evaluate the significance of fixed effects, we performed a Type III analysis of variance with Satterthwaite’s method for estimating degrees of freedom, as implemented in the lmerTest package in R. This method was selected due to its suitability for models with unbalanced designs and random effects, providing accurate and conservative inference for mixed models. Statistical significance was defined as *p* < 0.05. Post hoc pairwise comparisons between timepoints were conducted using estimated marginal means via the emmeans package in R.

## RESULTS

The primary factor influencing the release of cfDNA into the interstitium is thought to be cell death^21,23^. However, the temporal components of cfDNA release and its relationship to cell growth, cell death, and environmental effects such as drug exposure and the microenvironment remain underexplored. Here we set out to investigate the longitudinal dynamics of cfDNA release in monoculture, co-culture, and under the pressure of drug treatment using patient-derived xenograft (PDX) model systems.

### Differential growth and cfDNA fragmentation patterns in GBM43 and GBM12

GBM12 and GBM43 are well-characterized PDX models that retain their host’s original genomic profile, show a high degree of heterogeneity in culture, and form highly invasive tumors *in vivo*^25^. To assess cfDNA release over time, GBM12 and GBM43 cells were cultured over a five-day time course, during which cell counts, imaging, and supernatant-derived cfDNA measurements were collected at 24-hour intervals (**Figure 1A**). Morphologically, GBM12 cells form neurospheres whereas GBM43 cells grow as an adherent culture (**Figure 1B**). Over the course of five days, GBM43 cells demonstrated exponential growth with low levels of cell death (5-8%) across all time points (**Supplemental Figure 1A**). In contrast, GBM12 growth stalled during the first 48 hours, followed by rapid proliferation from 48-96 hours with a subsequent decline in expansion from 96-120 hours (**Supplemental Figure 1C**). These differences in growth dynamics were mirrored in cfDNA yield and fragmentation patterns. By 120 hours, GBM43 cells produced 61 ng of cfDNA, while GBM12 cells generated 1211 ng. This difference was confirmed using automated gel electrophoresis, which revealed clear nucleosomal cfDNA peaks in GBM12 but an absence of obvious peaks in GBM43 (**Supplemental Figure 1D-1E**). The appearance of this nucleosomal laddering pattern in GBM12 is suggestive of cell death by apoptosis. A correlational analysis of all triplicate data points for GBM43 revealed a moderate positive correlation between the number of dead cells and cfDNA yield. Surprisingly, a stronger and more statistically significant correlation was found between live cells and cfDNA yield in GBM43 time points suggesting an active secretion mechanism or high proliferative turnover characteristic of this cell line (**Figure 1C**). GBM12 exhibited a strong correlation between both live and dead cell counts and cfDNA yield across all time points; however, the correlation between live cells and cfDNA release was statistically significant, suggesting that in addition to cell death, active cellular processes or higher metabolic activity may also contribute to cfDNA production in this line (**Figure 1D**). Prior studies have shown that viable cancer cells can release cfDNA through pathways such as exosome-mediated export and amphisome-dependent trafficking, with actively proliferating or metabolically engaged cells contributing to cfDNA in supernatant^17,21,22,23^. These findings support the notion that cfDNA may serve not only as a marker of cell turnover but also as a dynamic indicator of tumor cell activity and intercellular communication.

**Figure 1.**
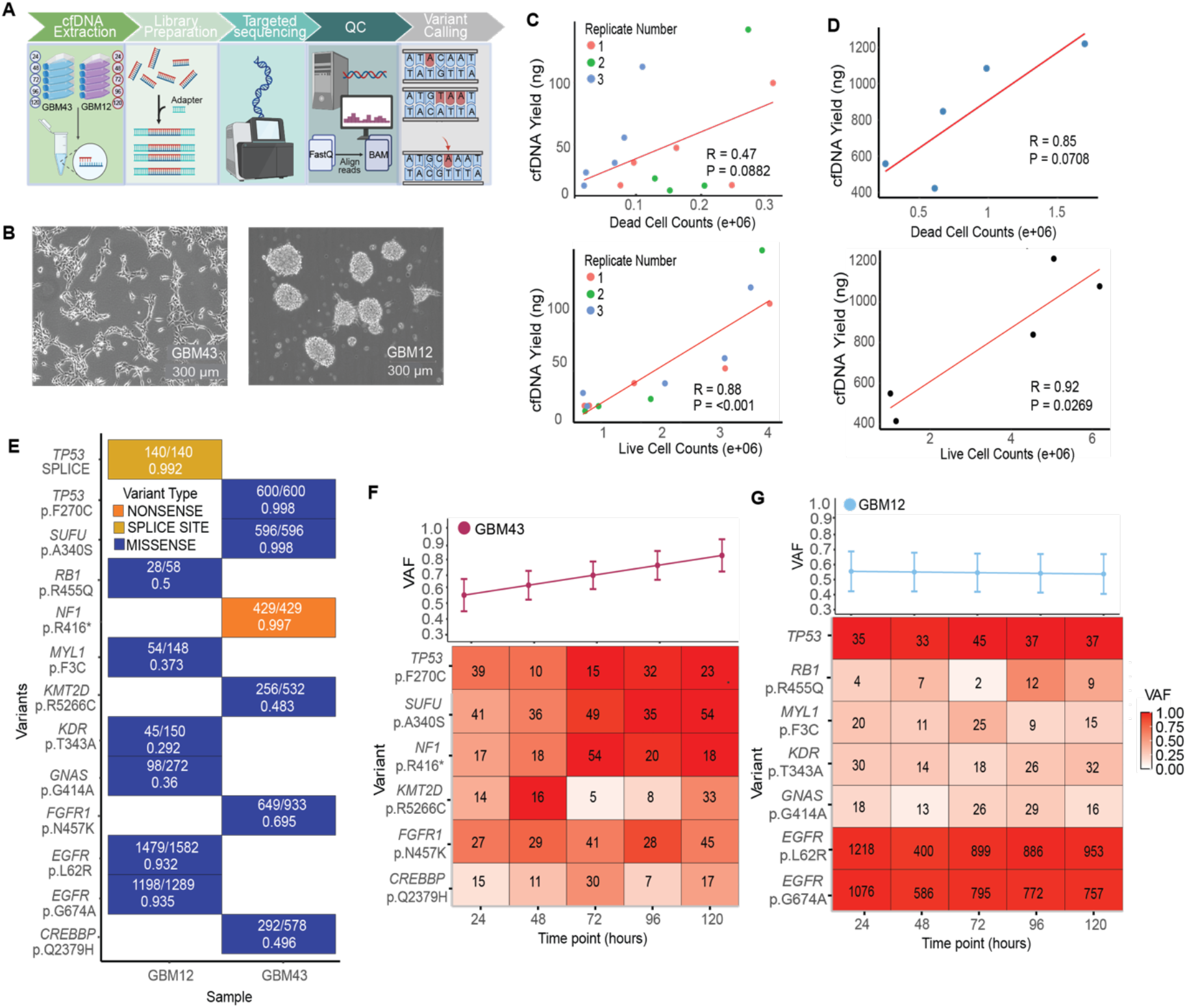
Study design and unperturbed monoculture baseline characteristics. **(A)** Schematic showing experimental timeline from cell culture to data analysis. **(B)** GBM43 (left) and GBM12 (right) cell morphology **(C)** Pearson correlation analysis showed a strong and statistically significant correlation with live cells and cfDNA yield in GBM43 triplicate experiments. **(D)** Pearson correlational analysis showed a strong correlation between both dead and live cells and cfDNA yield. **(E)** Genomic profiles of GBM43 and GBM12 gDNA. Each cell shows supporting reads / read depth and the VAF below. **(F)** Heatmap of VAF of GBM43 variants in cfDNA samples show statistically significant changes from 24 to 120 hours (Estimate = 0.0627 ± 0.0263, *t*(23) = 2.38, *p* = 0.026). **(G)** Heatmap of VAF for GBM12 variants in cfDNA samples did not show any statistically significant changes from 24 to 120 hours (Estimate = –0.0044 ± 0.0181, *t*(30) = –0.24, *p* = 0.8105).

### cfDNA Captures Tumor-Specific Mutations and Reveals Dynamic Variant Changes in GBM43 Cells

Targeted sequencing of genomic DNA (gDNA) from GBM43 and GBM12 revealed several unique variants. GBM43 gDNA contained five missense variants—*CREBBP* p.Q2379H, *FGFR1* p.N457K, *KMT2D* p.R5266C, *SUFU* p.A340S, and *TP53* p.F270C—along with one nonsense variant, *NF1* p.R416* (**Figure 1E**). In contrast, the mutational profile of GBM12 gDNA included six missense variants, and one splice site variant (**Figure 1E**). Notably, two *EGFR* missense variants (p.L62R and p.G674A) were identified at very high variant allele frequency (VAF) and at 8-to 10-fold increased coverage, indicative of a potential extra-chromosomal DNA (ecDNA) amplification. In glioblastoma, ecDNA is a well-established mechanism for oncogene amplification, particularly of *EGFR*, which enables high-level gene expression independent of linear chromosomal constraints^19,20^. All of the cell-line specific variants could also be observed in supernatant-derived cfDNA, demonstrating the potential of cfDNA to track the dynamics of tumor-specific variants in both model systems.

To evaluate changes in tumor-specific VAFs across serial timepoints in the GBM43 cell line, we performed a linear mixed-effects model with timepoint as a fixed effect and replicate as a random effect (**Figure 1F**). This model accounts for repeated measurements and inter-replicate variability. We observed a significant positive association between timepoint and VAF (*p* = 0.0259) in GBM43 cells, indicating that the VAF increased at a rate of 6.3% every 24 hours on average **(Figure 1F)**. Upon inspection of the individual variants, this increase appears most pronounced for 3/5 variants, including the variants in *TP53* and *NF1*, suggesting an ongoing adaptation to standard cell culture conditions (**Supplemental Figure 1F**). Applying the same statistical model to serial time points collected from GBM12, we observed a stable tumor-specific VAF across all cfDNA timepoints that was not statistically significant (*p* = 0.8105), indicating a steady tumor state (**Figure 1G**).

### GBM43 cells outcompete NHA cells in a co-culture system

To further explore cfDNA release in the presence of a tumor microenvironment, we co-cultured GBM43 cells with normal human astrocyte (NHA) cells (**Figure 2A**). Astrocytes, which are common in the glioblastoma tumor microenvironment, can transform into tumor-associated reactive astrocytes, enhancing tumor invasiveness and aggressiveness^18^. Baseline studies of the two cell types grown in monoculture showed that GBM43 cells grew exponentially, maintaining over 90% viability across all time points (**Figure 2B**, pink). NHA cells, in contrast, exhibited slower growth with comparable viability (**Figure 2B**, green). The differences in growth rates were also reflected in cfDNA yield. GBM43 cells exhibited a steady increase in cfDNA yield, rising from 6.1 ng at 24 hours to 61 ng at 120 hours (**Figure 2D**). This consistent increase likely reflects a combination of low levels of cell death and tumor-specific processes that facilitate the release of cfDNA, potentially through active secretion or turnover mechanisms associated with tumor growth.

**Figure 2.**
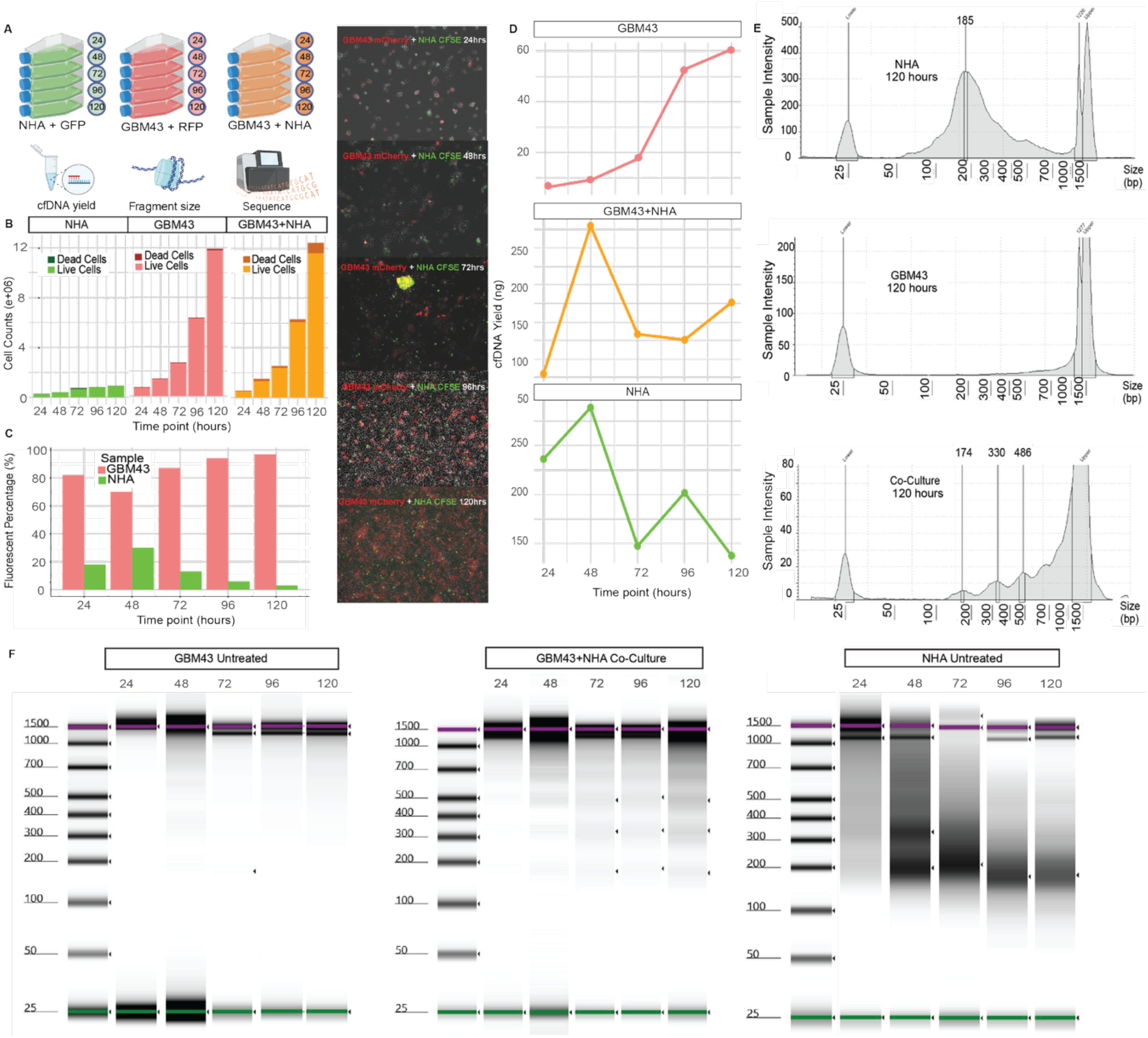
Competition of cell-free DNA release under co-culture conditions. **(A)** Schematic showing experimental set-up for GBM43 and NHA co-culture. **(B)** Difference in growth rates within experimental groups. **(C)** Distinct cell groups were found at varying levels at each time point in the co-culture using fluorescence. Images at each time point on the right. **(D)** cfDNA yield was highly variable within experimental groups. **(E)** Fragmentation patterns at 120 hours for co-culture differed from both individual cell lines. **(F)** Tapestation gel electrophoresis show differential fragmentation patterns between all timepoints in GBM43, co-culture and NHA.

In contrast, NHA cells demonstrated a decline in cfDNA yield over the same time period, decreasing from 212 ng at 24 hours to 113 ng at 120 hours despite comparable viability to GBM43 cells (**Figure 2D**). This decline suggests a rapid release of cfDNA at NHA culture inoculation and the subsequent decrease of cfDNA production per unit of time, despite consistent degradation of cfDNA.

The high cfDNA yield in NHA cells may reflect an early response to culture conditions, potentially driven by stress-induced cell death or active cfDNA shedding mechanisms. In contrast, GBM43 cells, which are characterized by their rapid proliferation and lower baseline cell turnover, showed much lower cfDNA release at early time points. In the co-culture system, overall cell growth was slightly higher than in the individual cultures (**Figure 2B**, orange). However, fluorescence quantification revealed that NHA cells were progressively outcompeted by GBM43 cells for nutrients and space. Over 24 to 120 hours, the proportion of NHA cells in the co-culture dropped from 50% to 19%, accompanied by increased NHA cell death (**Figure 2C**).

### Fragmentation pattern in co-culture showed unique patterns compared to monoculture

Fragmentation patterns varied distinctly across the different conditions. GBM43 cells exhibited no consistent fragmentation pattern, whereas NHA cells consistently produced mono-nucleosomal cfDNA, with fragments primarily around 185 bp, at all the time points (**Figure 2F**). In contrast, at 120 hours, the co-culture produced a unique cfDNA pattern with distinct peaks at 174 bp, 330 bp, and 486 bp. This pattern was distinct from the baseline fragmentation of either individual cell line (**Figure 2E**). The pattern in the co-culture might be indicative of a unique interaction between the tumor cells and the normal cells in a tumor microenvironment leading to differences in cfDNA release patterns compared to when the cell types are grown in isolation. This pattern may reflect a specific mode of cell death, or competition between the cell types in the co-culture, such as an interplay between apoptosis and necrosis that could result in a varied cfDNA release profile.

### VAF of co-culture cfDNA showed higher NHA cfDNA presence

Targeted sequencing of GBM43 and NHA gDNA showed several shared and unique variants (**Figure 3A**). Specifically, we identified *n* = 39 missense, *n* = 3 five-prime flank, and *n* = 1 splice site variant in the NHA gDNA. Comparatively, in GBM43 cells, we found *n* = 34 missense, *n* = 2 in-frame deletions, *n* = 2 five-prime flank and *n* = 1 nonsense variant. Amongst these, *n* = 25 missense variants were found to be unique to NHA, and *n* = 20 missense variants were unique to GBM43 (**Figure 3A**). Accordingly, there were *n* = 12 missense variants shared between GBM43 and NHA, representing common germline variants shared between the two models.

**Figure 3.**
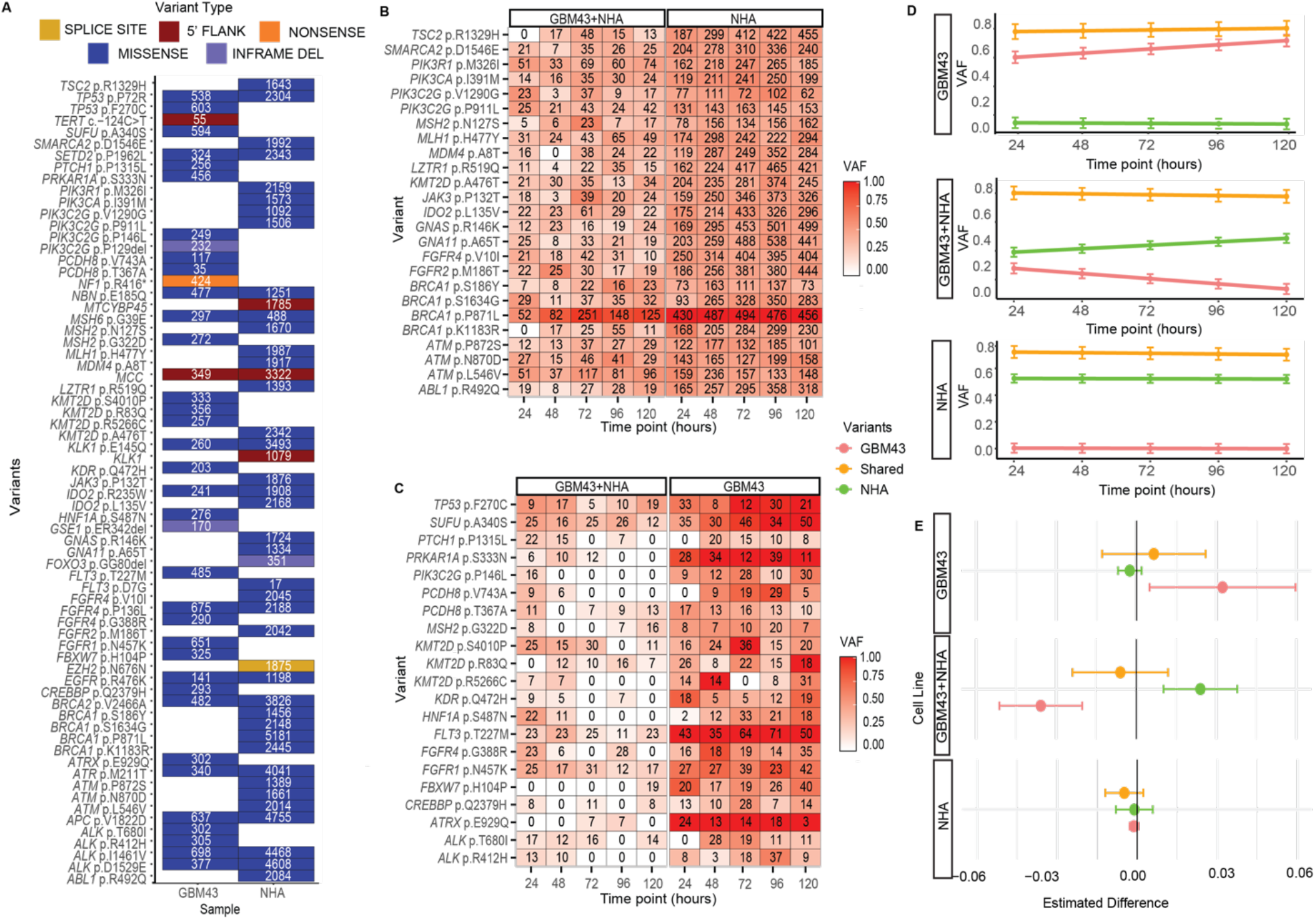
Longitudinal genomic analysis of cfDNA in a co-culture system. **(A)** NHA and GBM43 gDNA show several shared and unique variants. **(B)** Several NHA-specific missense variants were found in co-culture timepoints. **(C)** Several GBM43 unique missense variants were found in lower VAF in co-culture timepoints. **(D)** Co-culture showed a significant decrease in VAF from first to last time point in GBM43 unique variants (estimate = −0.0358, SE = 0.0078, df = 83, p < 0.001) and a significant increase in VAF from first to last time point in NHA unique variants (estimate = 0.0240, SE = 0.0070, df = 103, *p* = 0.000879). GBM43 unique variants showed a significant increase in GBM43 cfDNA (estimate = 0.03223, SE = 0.01372, df = 83, *p* = 0.0212). No other group–variant intersections demonstrated statistically significant changes in VAF. **(E)** Forest plot showing the estimated change in variant allele frequency (VAF) every 24 hours for each cell line within each variant group.

Utilizing the unique missense variants characteristic of each line, we were able to track temporal changes in VAF using serial cfDNA timepoints from mono- and co-culture experiments (**Figure 3B and 3C**). Turning again to a linear mixed-effects model to measure changes in VAF over time, we noted a stable average VAF of NHA-specific variants across all NHA monoculture cfDNA time points (*p* = 0.734). Conversely, GBM43-unique variants exhibited a statistically significant increase in VAF in cfDNA derived from a GBM43 monoculture (*p* = 0.0212), suggesting an ongoing evolutionary process in GBM43 cells that is absent in NHA cells. Intriguingly, in the co-culture experiment, GBM43-unique variants showed a decrease of 3.6% in VAF every 24 hours (*p* < 0.001). Correspondingly, NHA-unique variants exhibited an increase of 2.4% in VAF per 24-hour period in co-culture samples (*p* = 0.0008, **Figure 3D-E**). This inverse relationship was mirrored by cell count data, which revealed a progressive decline in NHA viability and a concurrent dominance of GBM43 cells over time. As NHA cells were increasingly outcompeted, fluorescence-based cell death measurements showed elevated NHA cell death, leading to greater cfDNA release from dying normal cells. This was reflected in both rising total cfDNA yield and a shift in variant representation toward NHA-specific VAFs. Fragmentation analysis further supported these findings, with co-culture cfDNA showing prominent nucleosomal laddering indicative of apoptosis. Conversely, the relative decrease in GBM43-specific VAF, despite continued proliferation, likely reflects reduced GBM43 cell death and less cfDNA release per cell. Together, these data illustrate how differential cell survival, proliferation, and death contribute to the evolving composition of cfDNA in a shared microenvironment.

### TMZ treatment affects cfDNA fragmentation and yield

Temozolomide (TMZ) is an oral alkylating agent and the standard-of-care chemotherapeutic used in glioblastoma treatment, typically administered alongside radiotherapy. It induces DNA damage by methylating guanine residues, leading to replication stress, cell cycle arrest, and ultimately apoptosis in rapidly dividing tumor cells^24^. To explore how TMZ-induced cell death impacts cfDNA release and fragmentation, we assessed the relationships between cell growth, cell death, and cfDNA dynamics in treated and untreated glioblastoma cultures in a similar time course experiment (**Figure 4A**). Treated cells exhibited minimal to no growth when compared to untreated cells, reflecting the efficacy of the 50% inhibitory concentration (IC50) of TMZ used in this experiment (**Supplemental Figure 2A-B**, red). As expected of TMZ-treated cells, the number of dead cells grew considerably over time (**Supplemental Figure 2B**, blue).

**Figure 4.**
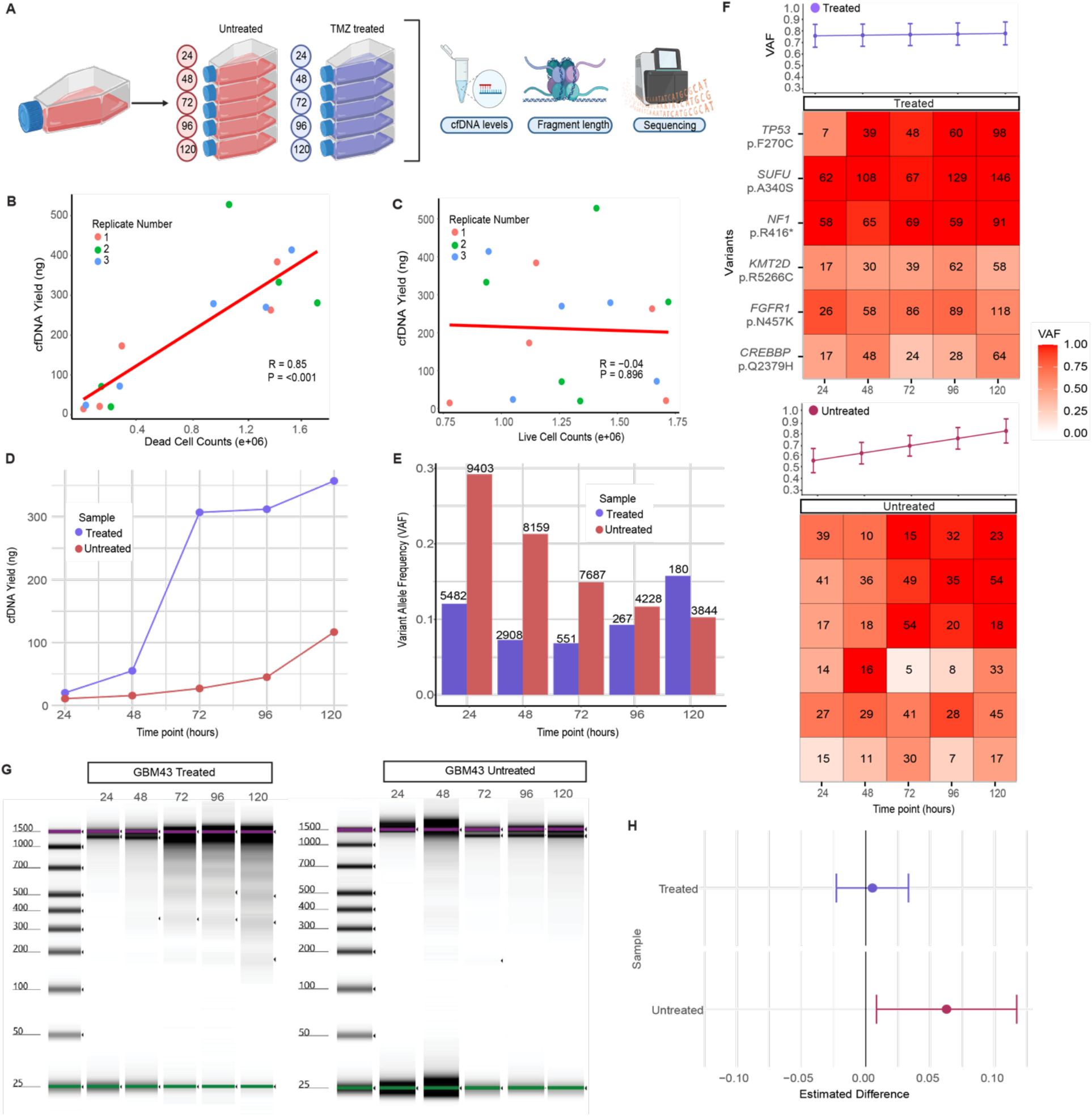
cfDNA release in response to TMZ treatment. **(A)** Schematic showing the set-up of GBM43 TMZ experiment. **(B)** Significant correlation between dead cells and cfDNA yield was found in treated triplicate data (p value calculated by Pearson) **(C)** Low correlation between live cells and cfDNA yield was found in treated triplicate data (p value calculated by Pearson). **(D)** Visible increase in cfDNA yield was seen primarily in the treated timepoints. **(E)** Overall VAF showed an increase in treated groups while the number of variants at each timepoint decreased (as indicated with the numbers at the top of the bars) while a decrease in VAF was associated with a decrease in the number of variants in untreated timepoints. **(F)** Forest plot showing estimated difference of the treated timepoints vs untreated timepoints. **(G)** Untreated cfDNA time points showed a significant increase in tumor-specific VAF first to last timepoint (estimate = 0.0627, SE = 0.0263, df = 23, *p* = 0.031) while treated time points show no statistical significance (estimate = 0.0052, SE = 0.0135, *p* = 0.705).

In untreated GBM43 cells, we observed a strong and statistically significant correlation between live cell counts and cfDNA release (**Figure 1C**), suggesting contributions from active secretion mechanisms. In contrast, TMZ-treated GBM43 cells exhibited a stronger correlation between cfDNA yield and the number of dead cells (**Figure 4B**), consistent with TMZ’s cytotoxic effects and induction of apoptosis. The untreated samples, which maintained low levels of cell death throughout the experiment, exhibited relatively stable and low cfDNA concentrations. In contrast, the treated samples showed a significant increase in cfDNA yield, consistent with the higher number of dead cells (**Figure 4D**). Notably, the treated condition at 72 hours exhibited a sharp rise in cfDNA concentration, corresponding to the dramatic increase in dead cells observed at this time point (**Supplementary Figure 2B**). While cfDNA has traditionally been associated with apoptosis and necrosis, our findings demonstrate that even within a single cell line, multiple cellular processes—including active release by viable cells—can contribute to cfDNA dynamics.

### Utility of cfDNA in capturing treatment selective pressure in GBM43

In treated GBM43 samples, the variant allele frequency (VAF) of gDNA-unique tumor variants remained consistently high across all timepoints (∼75%), whereas in untreated samples, VAF increased over time from approximately 55% at 24 hours to 80% at 120 hours (**Figure 4G**). Longitudinal analysis using a linear mixed-effects model revealed that treated samples did not show a statistically significant change in VAF over time (**Figure 4F**). These findings align with cfDNA yield trends, where increased cfDNA release was observed in response to TMZ treatment, supporting the utility of cfDNA as a biomarker for therapeutic response. To further investigate clonal dynamics, we calculated total variant allele frequencies by summing the VAFs of all detected variants at each time point and normalizing by total read depth. In the treated group, we observed a decrease in the number of detectable variants (noted above each bar) accompanied by an increase in aggregate VAF (**Figure 4E**, purple bars). In contrast, the untreated group exhibited decreases in both variant count and overall VAF (**Figure 34**, red bars). The significant increase in VAF in untreated samples reflects ongoing tumor evolution, while the stable VAFs in treated samples, despite reduced variant diversity, point to selective pressures imposed by therapy. This divergence underscores cfDNA’s utility in capturing distinct clonal trajectories under therapeutic versus natural progression conditions.

### cfDNA fragmentation patterns in response to TMZ treatment show clear apoptotic patterns

Apoptotic cfDNA fragments exhibit a characteristic laddering pattern due to nucleosomal DNA wrapping. Mononucleosomal fragments, containing approximately 167 bp of DNA, form the primary peak, with additional peaks corresponding to dinucleosomal (∼334 bp) and trinucleosomal (∼501 bp) fragments. In untreated samples, cfDNA fragmentation patterns were largely absent, even at 120 hours, which aligns with the low levels of cell death observed in this condition (**Figure 4G**). In contrast, treated samples displayed distinct nucleosomal peaks at 186 bp, 344 bp, and 491 bp, indicative of apoptotic cfDNA release (**Supplemental Figure 2C**). This apoptotic laddering pattern accentuates the mechanism of TMZ-induced cell death. Interestingly, although 3D GBM12 cultures exhibited similar levels of cell death, their cfDNA fragmentation patterns differed. GBM12 cfDNA displayed slightly longer fragment sizes compared to TMZ-treated GBM43, potentially reflecting differences in chromatin accessibility, nuclease activity, or cfDNA release mechanisms between the two GBM lines (**Supplemental Figure 2D**).

## DISCUSSION

Our study demonstrates the utility of cfDNA as a non-invasive biomarker for monitoring tumor dynamics, treatment response, and microenvironmental interactions in glioblastoma. Using in-vitro models, we tracked cfDNA yield, fragmentation patterns, and variant allele frequencies across experimental conditions including temozolomide (TMZ) treatment and tumor–normal co-culture systems. These findings provide insight into mechanisms of cfDNA release and highlight the translational potential of cfDNA-based monitoring in GBM.

The differences in cfDNA yield and fragmentation between GBM43 and GBM12 highlight the molecular and phenotypic heterogeneity of glioblastoma. GBM12 released markedly higher levels of cfDNA and exhibited well-defined nucleosomal fragmentation, indicative of apoptosis. In contrast, GBM43 displayed lower cfDNA yield and lacked apoptotic fragmentation patterns under baseline conditions. Interestingly, we observed a statistically significant correlation between cfDNA yield and live cells in untreated GBM43, suggesting that actively proliferating cells may also contribute to cfDNA release via non-lethal mechanisms such as exosome or amphisome pathways. These observations demonstrate that cfDNA reflects a broader range of biological processes beyond cell death and must be interpreted within the cellular context of each tumor model.

In co-culture experiments, we observed dynamic shifts in cfDNA composition and VAFs over time. While GBM43 cells ultimately outcompeted NHA cells in cell counts, the NHA-derived cfDNA increased steadily over time, driven by elevated NHA cell death in response to competitive stress. This was reflected in a corresponding rise in NHA-specific VAFs and cfDNA yield, as well as apoptotic fragmentation patterns resembling those seen in NHA monocultures. Conversely, GBM43-specific VAFs decreased, not due to a loss of tumor cells, but due to the relative reduction in cfDNA contribution from the tumor population. These findings illustrate how cfDNA can capture not only tumor burden but also competitive dynamics within the tumor microenvironment. The ability to distinguish cfDNA contributions from different cell types in co-culture was further supported by targeted sequencing of cell line-specific variants. Unique variants from GBM43 and NHA were reliably detected in their respective monocultures and successfully traced within mixed cfDNA pools from co-cultures, demonstrating the power of variant-informed deconvolution in heterogeneous settings. This has direct clinical relevance, where cfDNA often reflects contributions from both tumor and non-tumor sources.

Following TMZ treatment, GBM43 cells exhibited a clear increase in cfDNA yield accompanied by distinct nucleosomal peaks, consistent with apoptosis-mediated DNA fragmentation. VAFs of GBM43-specific variants also increased, paralleling cytotoxic effects and reinforcing cfDNA’s utility in measuring therapeutic response. These observations align with the known mechanism of TMZ as a DNA-damaging agent and validate cfDNA as a marker for treatment-induced cell death.

Overall, our findings highlight cfDNA as a sensitive and dynamic biomarker for glioblastoma biology. The ability to track tumor-specific variants, differentiate cell types in mixed environments, and correlate cfDNA profiles with cellular processes—including proliferation, death, and competition—provides a foundation for more precise, non-invasive disease monitoring. While our study was limited to in-vitro systems, the principles observed here have strong translational potential and should be validated in clinical settings, particularly for tracking minimal residual disease (MRD) and recurrence. Future work integrating cfDNA with additional liquid biopsy modalities, such as circulating tumor cells (CTCs) or extracellular vesicles (EVs), will be essential for building a comprehensive view of tumor evolution and treatment response.

With further validation, cfDNA holds promise for transforming GBM management through real-time, minimally invasive monitoring of tumor burden, therapeutic efficacy, and resistance dynamics.

## ACKNOWLEDGEMENTS

We would like to thank members of the Barthel and Berens groups for valuable discussions that have improved the rigor of our experimentation and interpretation of the results.

## AUTHORSHIP STATEMENT

SM and FB contributed to study conception and design. SM, HN performed all cell culture experiments and data collection. SM and MM conducted sample processing and sequencing. SM and MK completed analysis. SM, FB and MB assisted in revisions and interpretation of results. NT and MB provided cell lines for experiments. All authors read and approved the final manuscript.

## DATA AVAILABILITY

The datasets generated and/or analyzed during the current study are available from the corresponding author upon reasonable request. Targeted sequencing data will be made available in SRA under BioProject ID PRJNA1287894 upon publication.

