## Supplemental figures for "Cell-Free DNA Sequencing Uncovers the Longitudinal Consequences of Temozolomide Treatment and Host Co-Culture in Glioblastoma"

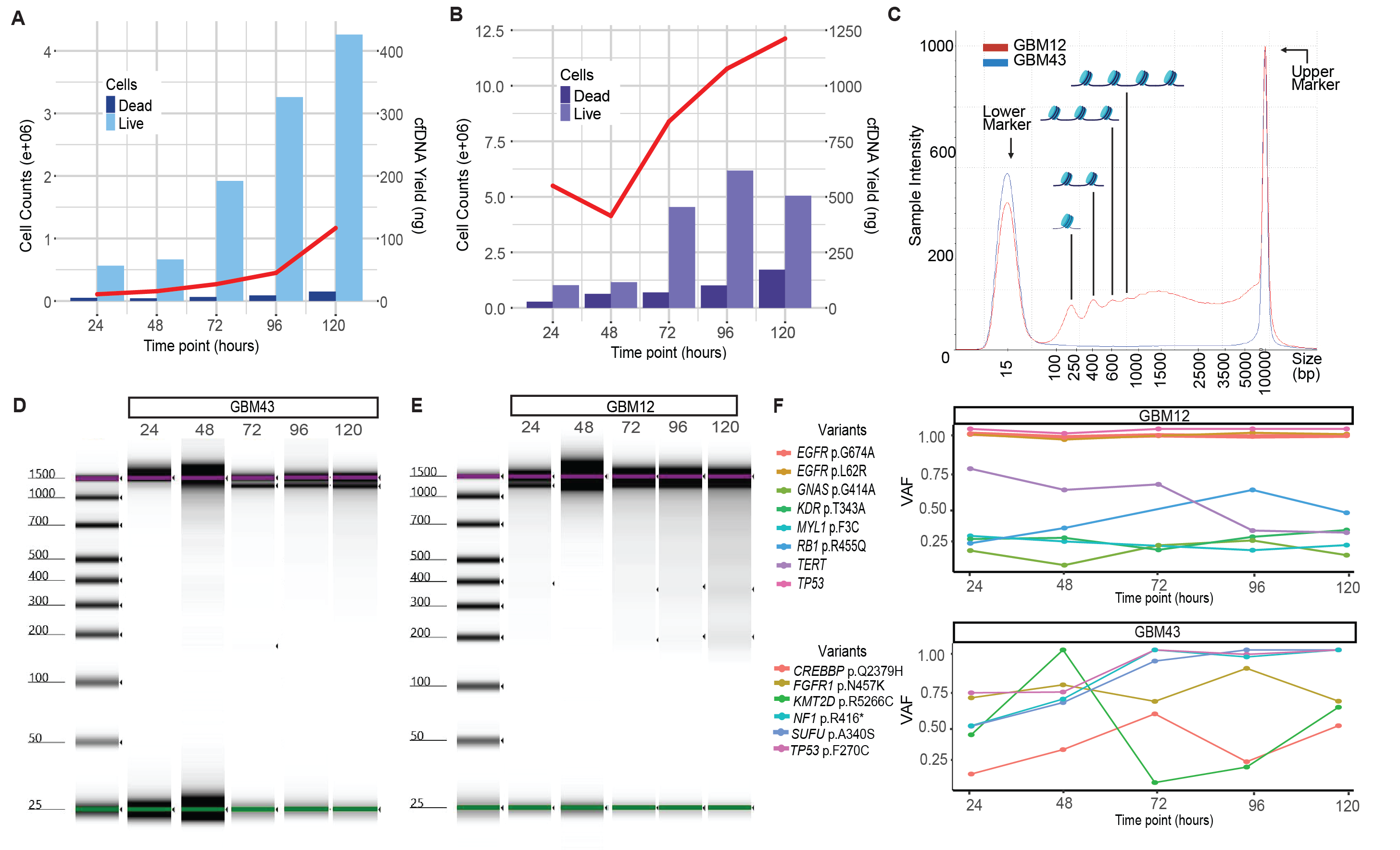

**Supplemental Figure 1**. **(A)** Rapid growth rate of GBM43 cells associated with steady increase in cfDNA yield. **(B)** Spike in growth rate at 72 hours for GBM12 cells cause varying levels of cfDNA throughout the time course. Each cell culture experiment was run in triplicate and cell counts (live and dead) and cfDNA yield was averaged together across all data points to create the plots seen above. **(C)** Size analysis reveals differences in average fragment size for each nucleosomal peak between GBM cell lines. **(D-E)** Tapestation gels for all timepoints in GBM43 and GBM12. **(F)** Individual VAF over time for GBM12 unique variants. Individual VAF over time for GBM43 unique variants.

**Supplemental Figure 2. (A)** The concentration of TMZ delivered to GBM43 cells was determined via DDR assay. **(B)** Increased cell counts for untreated cells shows GBM43 exponential growth rate. **(C)** Presence of nucleosomal-wrapped cfDNA can be detected in TMZ-treated timepoints. **(D)** GBM12 shows longer fragments at each peak indicating a difference in release mechanism between the two cell lines

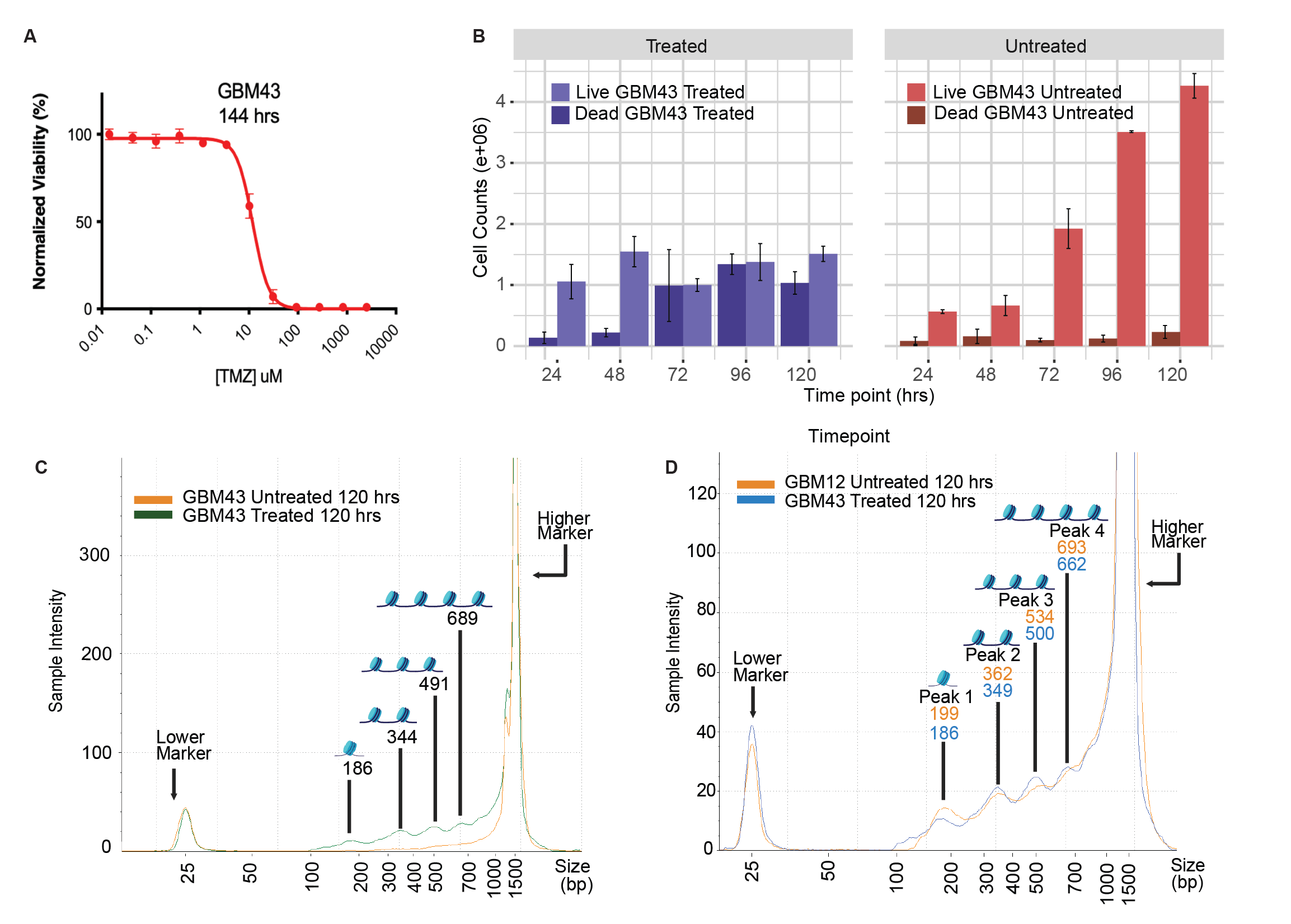

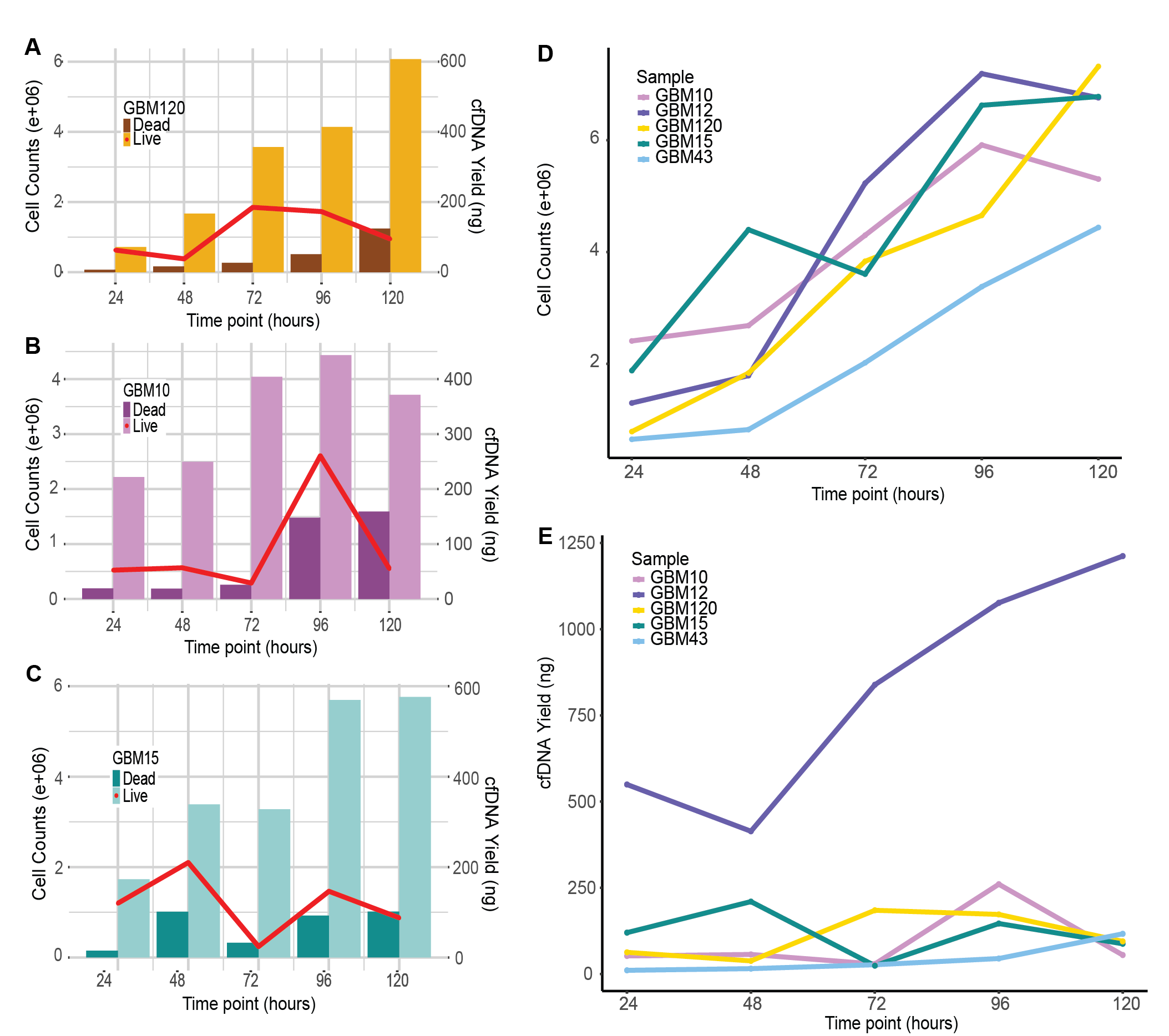

**Supplemental Figure 3. (A-C)** Cell counts and cfDNA yield across 24-120 hours for GBM120, 10 and 15 respectively. **(D)** GBM12 shows the highest cfDNA yield amongst cell lines. **(E)** All neurosphere lines show similar rates of growth.

**Supplemental Figure 4. (A-B)** Growth and cfDNA yield for GBM43 untreated triplicate cell culture. **(C-D)** Growth and cfDNA yield for GBM43 treated triplicate cell culture.

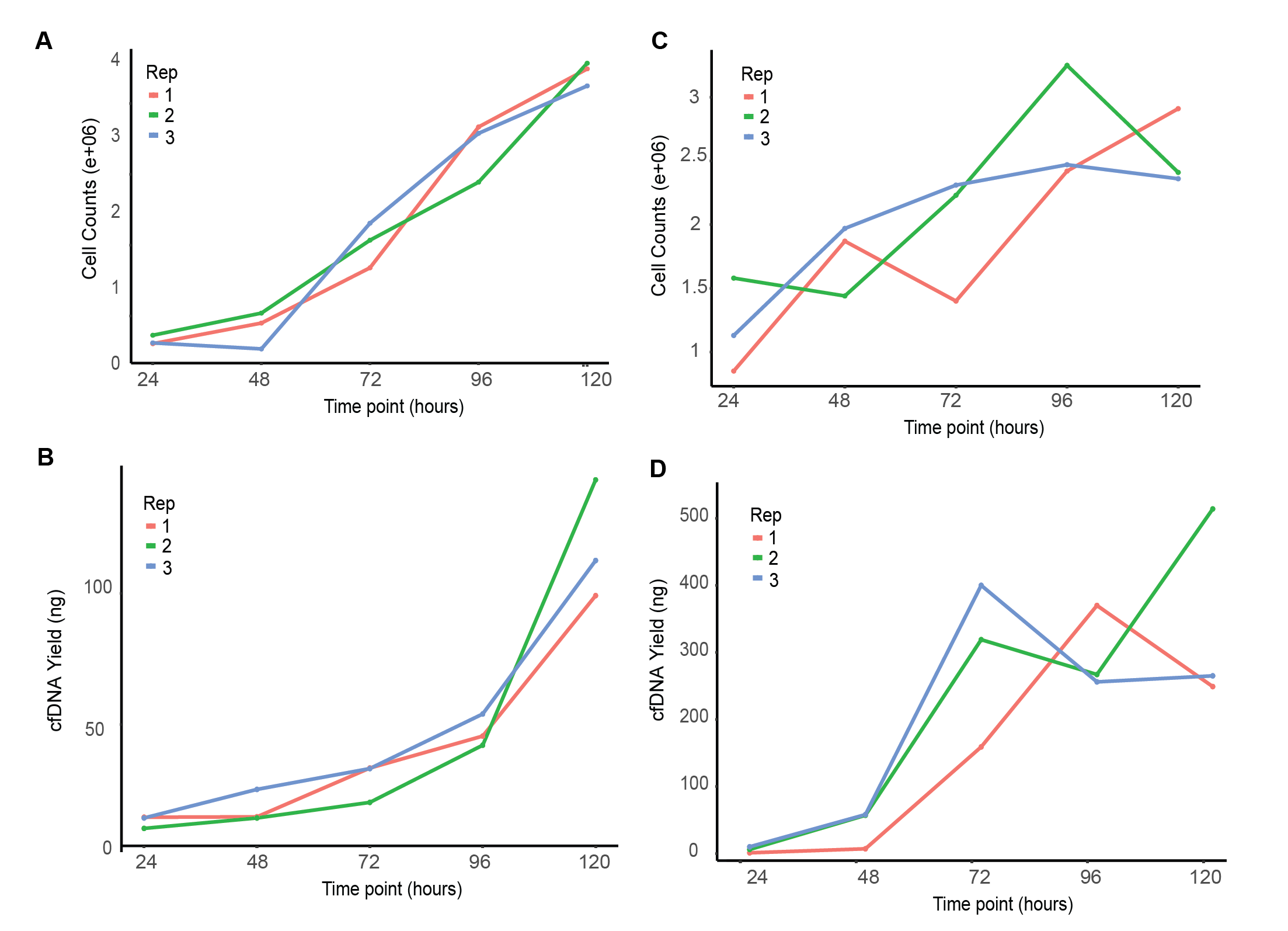

**Supplemental Figure 5. (A)** Individual VAF for GBM43 unique variants. **(B)** Individual VAF for NHA unique variants. **(C)** Individual VAF for NHA and GBM43 shared variants.

ssssssx

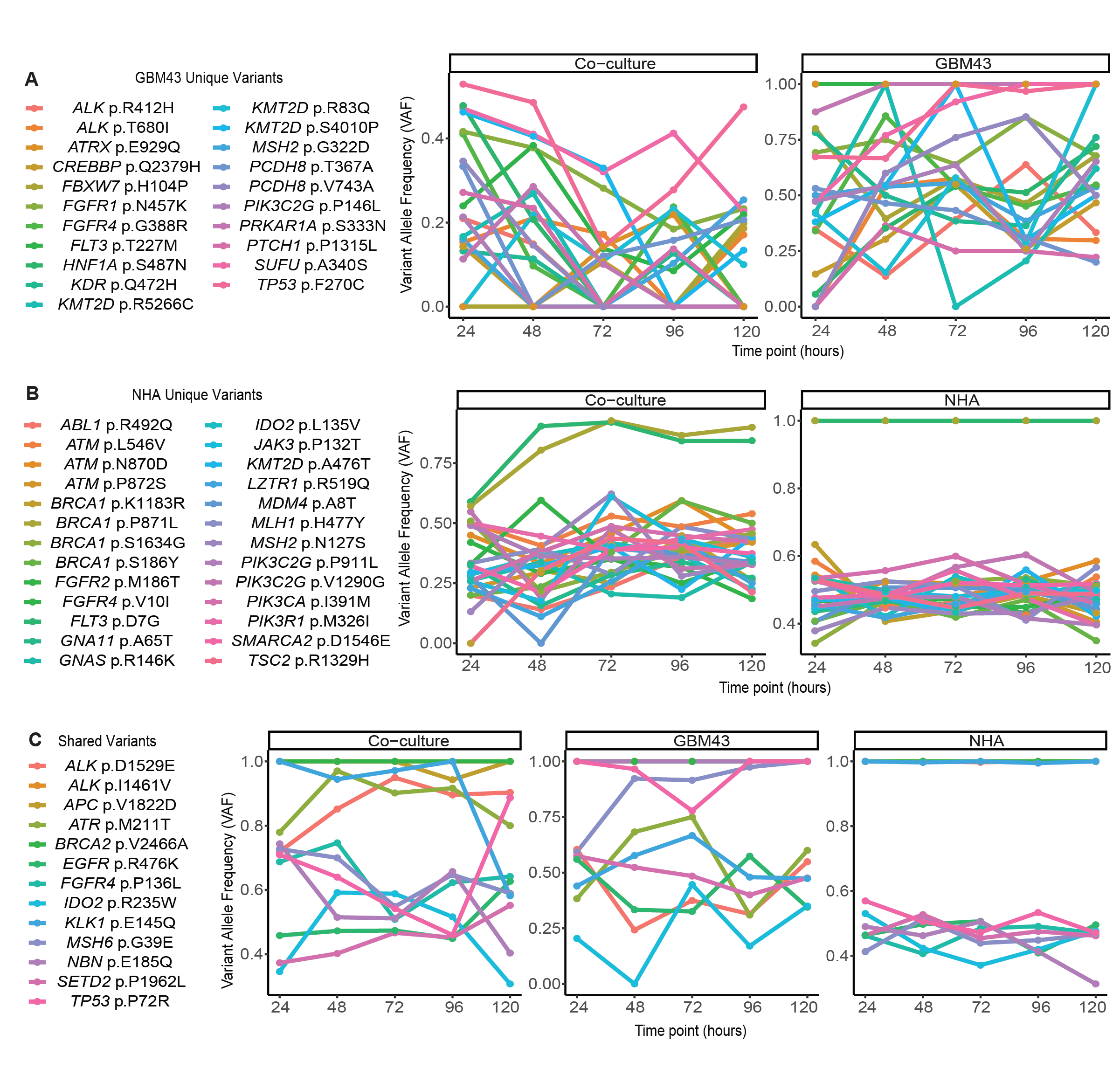
